# Large-scale atmospheric circulation cells regulate migratory dynamics of the iconic monarch butterfly

**DOI:** 10.64898/2026.07.16.738965

**Authors:** Naresh Neupane, Leslie Ries, Robert P. Guralnick, Elise A. Larsen

**Affiliations:** Department of Biology, Georgetown University; Florida Museum of Natural History, University of Florida, Gainesville, FL 32611

## Abstract

Understanding the impacts of climate on migratory dynamics is a priority for global change research, especially as phenological mismatches have been identified as a primary concern for long-distance migration. The eastern monarch (*Danaus plexippus*) is a flagship species for insect migration, but there has been surprisingly little research on their spring and summer migratory timing and mismatches with their key breeding resources. We show that spring, but not summer, phenological mismatch with their milkweed host plants has the potential to be a limiting factor in the eastern monarch’s yearly population growth. To better understand arrival at their breeding grounds, we developed models specifically to tease apart the factors that push (from departure grounds), pull (towards arrival grounds), or facilitate flow (in the flight corridor) during migration. We found wind (flow) dynamics to be the most important factor associated with arrival timing, especially in spring. Further, we show how spring and summer migratory flyways coincide with the large-scale Hadley and Ferrel atmospheric cells respectively, and how their dynamics appear to be essential for understanding monarch migratory timing.

## Introduction

Winged aerial migrations are among nature’s most striking phenomena, transporting biomass and nutrients across vast distances and creating important teleconnections among ecosystems^1–4^. Migratory species have evolved the ability to track peak resources across locations often separated by thousands of kilometers^5,6^. But this strategy comes with energetic costs and cumulative stressors, placing migrants at elevated risk of decline^7,8^. Differentially shifting environmental conditions across this migratory cycle has emerged as one of the most concerning ecological impacts of climate change^5^, particularly because migrants often rely on proxy cues to anticipate distant resource availability^9^. Decoupling of these cues can result in phenological mismatches, sometimes driving demographic declines^10,11^ and cascading community effects^12^.

Studies of migratory phenology most commonly correlate arrival timing with conditions on arrival grounds, which allows assessment of potential mismatch with key resources^13^. However, distant resources are unlikely to be local cues that initiate migration; instead, spatiotemporal correlates in large-scale environmental conditions can offer proxy information^13–15^. Examining conditions at departure sites^16^ and throughout migratory corridors^17^ is therefore critical to understand the proximate factors driving observed patterns in timing. One challenge for comprehensive research on migration across a full annual cycle is that, for most taxa, we have limited knowledge of their migratory connectivity — the degree of convergence and/or segregation among individuals in a single migrating population across different seasonal ranges^18^. In other words, during multiple steps in a cycle we don’t know how cohesive the individuals within a population are, erecting a barrier to connecting environmental conditions across the cycle to population performance.

Another major obstacle to a general understanding of migratory dynamics is that research has disproportionately focused on vertebrates, despite insect migrations vastly exceeding them in species richness, abundance, and biomass^1,4^. Insects are a particularly important group, not only because they are experiencing global declines^19^, but because they comprise roughly 62% of described animal species and underpin most terrestrial food webs^20^. Unlike vertebrates, most round-trip insect migrants have multi-generational cycles in which individuals returning at the end of a full cycle are the descendents of those that originally left^4^. Similar to vertebrates, lack of information on a population’s migratory connectivity remains a primary obstacle for disentangling underlying mechanisms across the full migratory cycle for most species^21^.

Here we focus on the eastern population of the iconic migratory monarch butterfly (*Danaus plexippus*) to identify key environmental drivers of their spring and summer arrival timing. The eastern monarch is the best-studied, non-pest insect migrant^4^ and comprises a largely panmictic population (Fig. 1), effectively removing the obstacle of tracking migratory connectivity. Our first goal was to estimate arrival timing and host plant emergence to assess potential phenological mismatches. Our second was to implement a generalizable modeling framework to be able to tease apart factors governing migratory dynamics across an annual cycle. Our approach partitions candidate drivers into push, pull, and flow (hereafter, PPF) factors (Fig. 1). Push factors induce departure, and these could be a change in local conditions^16^, or the appearance of proxy cues. These cues include factors such as day length and sun position^14,22^ as well as atmospheric pressure, which can provide short- and long-range forecasts of favorable migratory conditions^23,24^. In contrast, pull factors refer to conditions at or near arrival grounds that influence movement direction and/or flight speed through the corridor.

**Figure 1.**
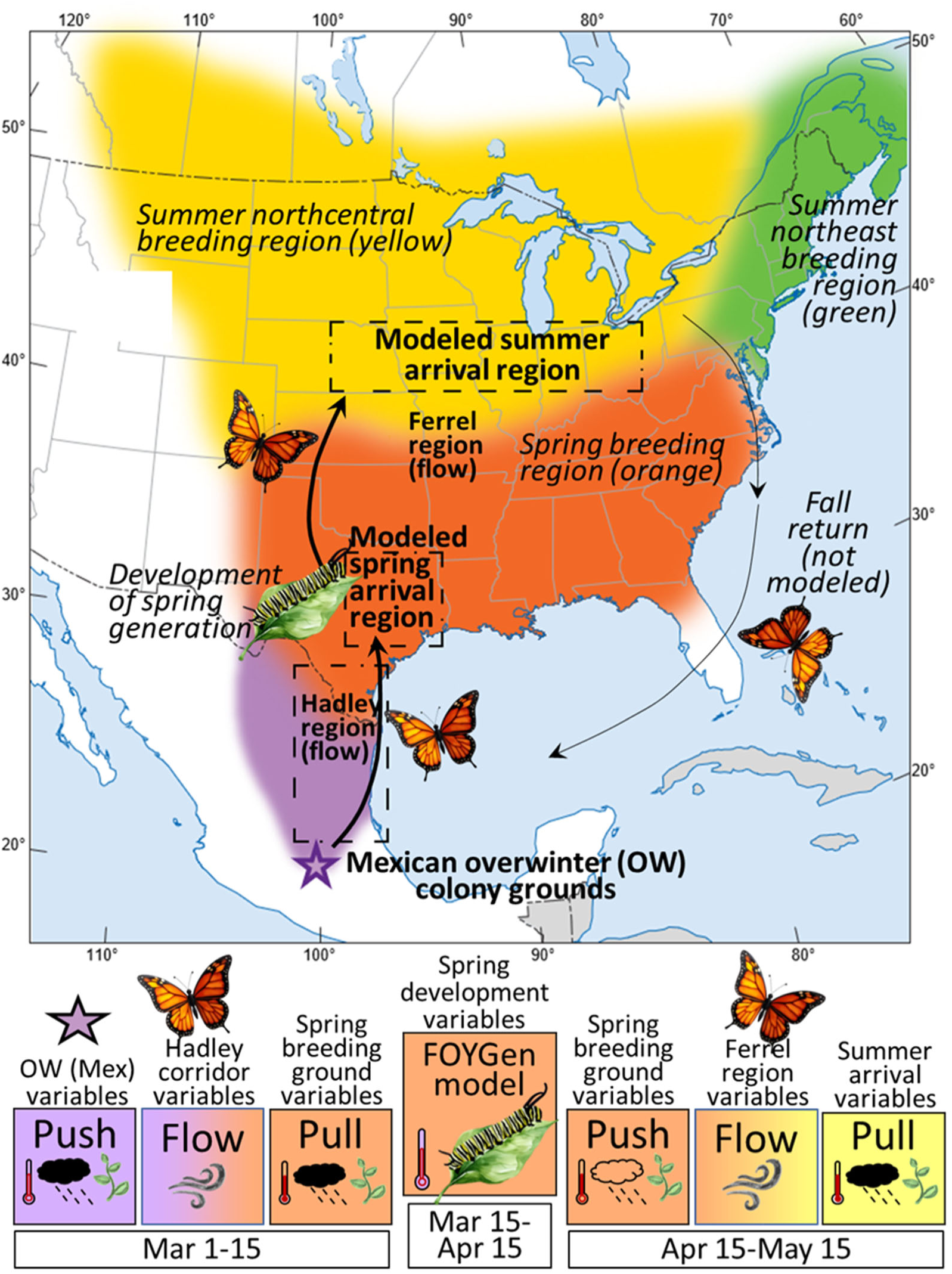
We model the phenology of Monarchs in 3 phases: arrival to the spring region, bridged by the development of the spring first-of-year (FOY) generation, and the subsequent flight to the northcentral summer breeding grounds. Arrival and flight models consider push factors in the departure region, pull factors in the arrival region, and large-scale weather patterns and nectar availability representing flow factors in between. Models are parameterized using Journey North (JN) sightings in the (all parameters are detailed in Table S1). *Adapted from Oberhauser et al*^61^.

Flow factors include atmospheric and ecological conditions spanning the migratory ranges, such as temperature, prevailing winds, or resources that facilitate movement^13,17,23^. Many aerial migrants are known to exploit tailwinds at different altitudes to speed flight trajectories, a phenomenon documented in birds^25^, bats^26,27^, and insects^28,29^, including monarchs^30^. These flow dynamics are often shaped by large-scale atmospheric circulation cells^31–34^ including the Hadley and Ferrel cells (Box 1, Panel A). In the context of these cells, the eastern monarch is particularly interesting because its spring and summer migratory phases show a particularly strong coincidence with the Hadley and Ferrel cells respectively (Fig. 1, Box 1). Specifically, the ascending branch (upward winds) of the Hadley cell is coincident with the monarch overwinter colony sites in Mexico (Box 1, Panel B). After winds ascend, they blow northward before descending within the latitudinal band where the majority of migrating monarchs are observed to arrive in spring. Suitable flight temperatures for monarchs across this vertical profile may also be an important flow factor because they will tend to push individuals down to the ground towards the location of the monarch’s typical arrival zone (Box 1, Panel B).

The eastern monarch population has experienced ongoing declines since at least the 1970s^35^ and is currently under consideration for threatened status under the U.S. Endangered Species Act^36,37^. Deforestation, host plant declines, pesticide use, and climate change are all implicated^35^. There has been extensive research on factors related to the monarch’s autumn migration, but evidence linking it to population declines has been weak^35^. In contrast, the potential role of alterations in spring and summer migratory timing and/or resource mismatch have received little attention, even though this is when the vast majority of reproduction occurs (reproduction does occur in autumn, but it is considered aberrant^38^). One study showed that there is currently no mismatch between monarchs and their hostplants at the beginning of summer, and monarch arrival would need to advance two weeks before demographic declines might become a concern during summer^39^. We could find no studies on phenological mismatch in spring, and this is likely due to the low density of monarchs on the landscape, which makes them difficult to study^40^. Yet spring recruitment is posited to be the most limiting factor in yearly population growth based on research partitioning environmental drivers of population growth across their full cycle^35,41^ This lack of research on breeding phenology, especially in spring, and its links to demography of eastern monarchs represents a significant gap in our knowledge. The first step to filling that gap is to understand the factors governing phenological timing and to measure the degree of mismatch currently occurring.

Our primary goals were to assess possible temporal mismatches between monarchs and milkweed and to model the factors governing the monarch’s migratory arrival timing. We focused on their central flight corridor (Fig. 1) because this is where a disproportionate number of arrivals at overwintering sites in Mexico have been shown to originate^35^. After quantifying overlap of milkweed emergence and monarch arrivals across their entire breeding range, we used our PPF framework to parse proximate environmental drivers during spring and summer migration (see Methods). We used a linear model to partition the relationship between different PPF environmental conditions and interannual variability in spring arrival timing. We then introduced a pause in the migratory progress to allow for development of the first-of-the-year (FOY) generation. This bridges the population’s movement from the spring to summer breeding ranges (Fig. 1), a phase that has not been incorporated into most predictive models of insect migration timing, but see^42^. Here, we used a deterministic model based on host plant emergence and monarch developmental temperature thresholds measured in a laboratory setting^43^. For the final phase, we again use a linear model to relate PPF factors to arrival timing at their summer breeding range in the Midwest (Fig. 1). We used 20 years (2001–2020) of phenology data collected from volunteer observers on the online platform, Journey North (JN)^44,45^, to estimate phenology metrics for monarch arrival and milkweed emergence (see Methods). Our initial hypothesis was that migratory timing will be rooted in large-scale atmospheric cells because of our *a priori* observation that the overwinter, spring, and summer ranges align with the positions of the Hadley and Ferrel cells (Box 1, Panel A). All data and code are available on github^46^.

## Results and Discussion

We show substantial overlap of spring monarch arrival with the beginning of milkweed emergence in TX, which occurs in March before a steep rise in April greenup (Fig. 2a). This suggests that monarchs arrive on their spring breeding grounds with smaller, sparser host plants relative to later in the season. In contrast, we show that milkweed emergence on the summer breeding range takes place a comfortable 2-3 weeks before most monarchs arrive (Fig. 2), similar to on-the-ground field surveys^39^. This substantial shift in phenological overlap begins at ∼34॰N (Fig. 2b) well to the south of their summer breeding grounds and this makes sense given the approximate one-month pause for development of the FOY generation (Fig. 1). We found that monarchs average arrival date rarely occurred before the average emergence date (this happened in only two years, by one and four days). However, even if mean arrival and emergence dates are the same, herbaceous plants are smaller and less dense at the start of their emergence period, suggesting that milkweed biomass will be low compared to later in the season. Further, milkweed is less dense overall in the south compared to northern breeding ranges^47^, Taken together, these two patterns suggest that oviposition sites for arriving monarchs in spring are likely to be much more limiting than in summer. This could be a crucial dynamic to consider because research has shown that during the early part of the season, monarch survival is most limited by egg density on individual host plants^48,49^. Females have even been shown to choose lower quality plants to avoid egg loading that could lead to starvation of emerging caterpillars^48^. This result is particularly important because spring conditions in eastern Texas have consistently been shown to be the most important factor predicting how large the population is able to grow each summer^35,41^.

**Fig 2:**
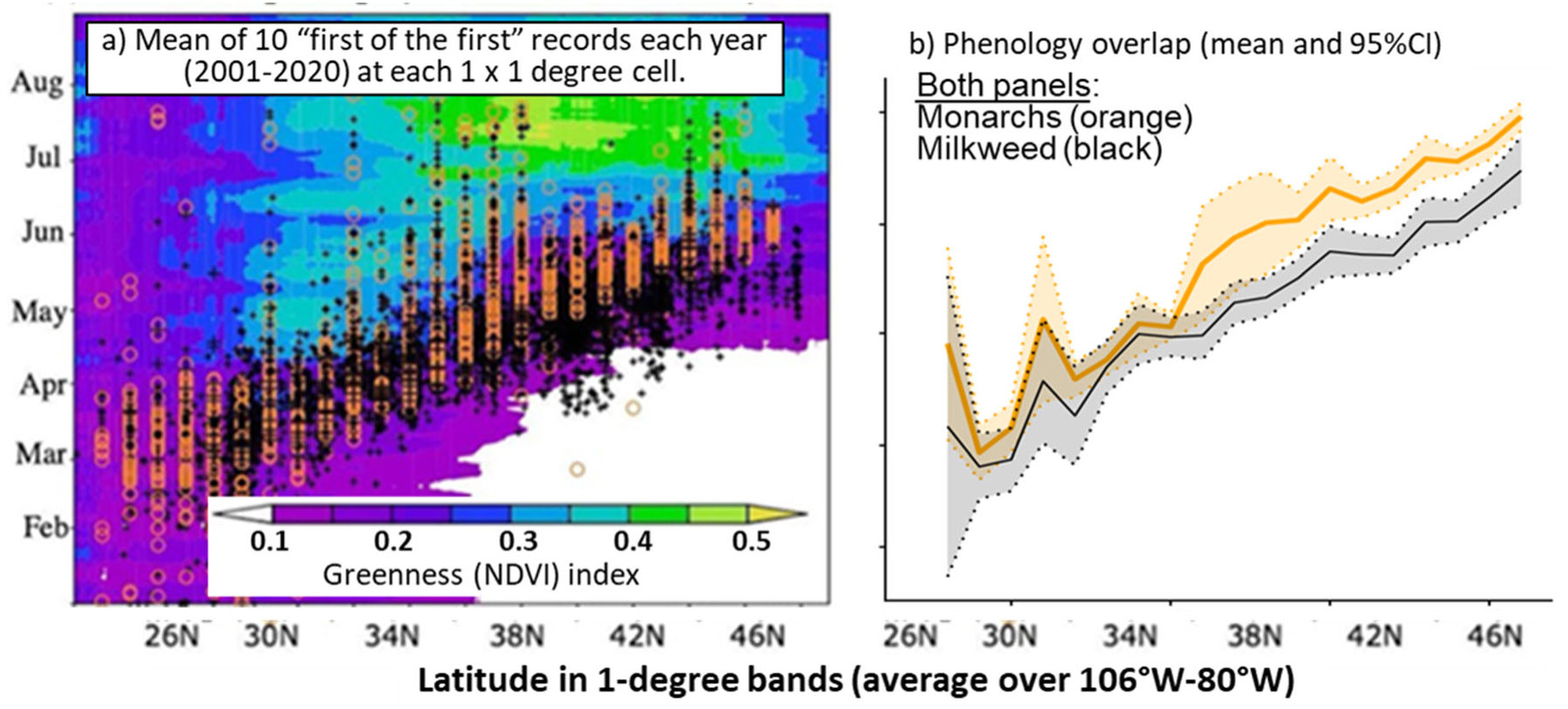
Phenological overlap of the first-reported sightings of monarch arrival and milkweed emergence from the Journey North platform where observers report their own first sightings each year. (a) Orange circles and black crosses indicate, respectively, 2001-2020 annual means of the days for the first ten observations for first monarch and milkweed sightings at 1-degree latitude resolution, pooled across 106°W-80°W and overlaid on a greenness (NDVI) index. (b) Across-year mean and 95% confidence intervals for each latitudinal band based on modeled responses.

Flow conditions were identified as the primary environmental factor explaining yearly variability in arrival timing, and this was especially true in spring (Table 1). During spring and summer migration, winds were blowing consistently up and north on departure grounds, then northward in the flyway (Table S2). Both spring and summer arrival timing were predicted to be earlier when temperatures were uniformly warmer across the migratory region and when wind was blowing more strongly northward in the migratory corridor (Table 2). In summer, the model that included all PPF phases (hereafter, all-combined) clearly outperformed the next best model (ΔAIC of 9.2) and included greenup timing as a push variable and precipitation as a pull variable (Table 1b). For spring arrival, the flow-only model had the lowest AIC score and included both wind and temperature variables (Table 1a). However, the all-combined was also indicated as a top model for spring (ΔAIC of 1.6), although there was little improvement in model performance (Table 1a). In spring, the all-combined model tied with the top-chosen PPF model. This model identified surface greenness in Texas (a proxy for milkweed and nectar availability) as a pull variable (Table 1a). This suggests that the presence of milkweed resources on spring breeding grounds could be a draw for monarchs. Yet it is important to note that milkweed is also available around the Mexican overwintering colonies to the north and south ^50^ suggesting that even if the presence of milkweed is a draw, that wouldn’t necessarily compel monarchs to fly northward. Overall, the influence of flow factors was more predictive in spring compared to summer (R^2^ = 0.76 and 0.57 respectively, Table 1), but this is not surprising because prevailing winds slow substantially and are more variable when approaching the Midwest arrival zone (Box 1, Panel C).

**Table 1.** Models predicting spring (phase 1) and summer (phase 3) arrival timing are bridged by a mechanistic model (phase 2) of development time of the FOY (first-of-the-year) generation. The summer arrival model propagates error across all three phases because it is based on estimated (not observed) values from the previous two phases. Selection among the candidate model concepts occurred in two stages. First, each concept model was implemented using all candidate variables categorized as push, pull, or flow (detailed in Table S1) and the variables chosen for each top concept model are shown for spring (a) and summer (b). Then, the best concept model was chosen among candidates using AIC (in bold, final parameter estimates shown in Table 2). All models included latitude and number of observations as an effort offset and are compared to a null model (see methods). Temperature (T) and wind (W) values were highly correlated across the entire flight zone so were transformed using principal components (PC) resulting in T.PC and W.PC components respectively (Table S2). T.PC loadings cannot be easily parsed into push, pull and flow, so they are allowed as variables for any component model, but grayed out for components that were not the focus of that model concept. The other push and pull components were precipitation (P), greenness (NDVI) and greenup (GrUp) in departure and arrival zones (Table S1). Model predictions and observations for each phase are shown in Fig. 3.

| <u>Candidate</u><br><u>model concepts</u> | <u>Push (MX)</u> | <u>Pull (TX)</u> | <u>PCA "Flow only"</u> | <u>AIC</u> | <u>Adj. R2</u> | <u>RMSE</u><br><u>(+/- days)</u> | <u>VIF</u> |
| --- | --- | --- | --- | --- | --- | --- | --- |
| **Flow (PCs only)** | *T.PC1, T.PC2* | *T.PC1, T.PC2* | **W.PC1, W.PC3,**<br>**W.PC5, T.PC1, T.PC2** | **163.5** | **0.76** | **3.3** | **2.5** |
| **All Combined** | **P, T.PC1, T.PC2** | **NDVI, GrUp,**<br>**T.PC1, T.PC2** | **W.PC1, W.PC3,**<br>**W.PC4, T.PC1, T.PC2** | **165.1** | **0.76** | **3.2** | **4.9** |
| Push-Pull only | P, T.PC1, T.PC2 | NDVI, GrUp,<br>T.PC1, T.PC2 | *T.PC1, T.PC2* | 170.9 | 0.73 | 3.5 | 2.0 |
| Push only | T.MX, P.MX | Not considered | Not considered | 188.8 | 0.62 | 4.3 | 1.3 |
| Pull only | Not considered | T.TX, P.TX | Not considered | 201.3 | 0.53 | 4.7 | 1.0 |
| Null | Not considered | Not considered | Not considered | 219.9 | 0.35 | 5.7 | NA |

| <u>Candidate</u><br><u>model concepts</u> | <u>Push (TX)</u> | <u>Pull (MW)</u> | <u>PCA "Flow only"</u> | <u>AIC</u> | <u>Adj. R2</u> | <u>RMSE</u><br><u>(+/- days)</u> | <u>VIF</u> |
| --- | --- | --- | --- | --- | --- | --- | --- |
| **All Combined\*** | **GrUp** | **P.MW** | **T.PC1 + W.PC2 +**<br>**W.PC3 + W.PC4 +**<br>**W.PC6** | **141** | **0.57** | **4.7** | **3.4** |
| Flow Only* | *T.PC1* | *T.PC1* | *T.PC1+W.PC2 +*<br>*W.PC3 + W.PC4 +*<br>*W.PC6* | 150.2 | 0.43 | 5.6 | 1.9 |
| Push-Pull only* | T.PC1, P, NDVI | T.PC1, P, NDVI,<br>GrUp | *T.PC1* | 150.5 | 0.42 | 5.6 | 1.5 |
| Pull only* | Not considered | T.MW, P, NDVI,<br>GrUp | Not considered | 156.4 | 0.30 | 6.4 | 1.4 |
| Push only | P, NDVI | Not considered | Not considered | 160.3 | 0.21 | 6.9 | 1.1 |
| Null | Not considered | Not considered | Not considered | 164.7 | 0.06 | 7.8 | 0 |

\*Chosen variables include the slope variable correcting estimates from phase 2 departure and flight times.

**Table 2:** Parameter estimates for the top models for spring (a) and summer (b) arrival timing chosen by AIC (Table 1). Details about all parameters included in each model are shown in Table S1t. Based on the PC loadings and our interpretations (Table S2), we provide our interpretations of each parameter retained in the model relative to its impact on arrival timing.

**a) Parameter estimates selected for the "Flow only" top concept model (Table 1a) for spring arrival**
| <u>Top model parameters</u> | <u>Estimate</u> | <u>Std. Error</u> | <u>t-value</u> | <u>Pr(>\|t\|)</u> | <u>Parameter interpretation</u> |
| --- | --- | --- | --- | --- | --- |
| (Intercept) | 68.75 | 0.79 | 86.92 | 5.6E-58 |  |
| Wind PC1 | -0.93 | 0.43 | -2.19 | 3.3E-02 | Stronger northward winds in the migratory corridor and more upward winds in TX |
| Wind PC3 | -3.09 | 0.77 | -3.99 | 2.1E-04 | Even stronger northward winds in corridor and TX, but relatively weaker upward winds in MX |
| Wind PC5 | -2.91 | 1.54 | -1.89 | 6.4E-02 | Even stronger northward winds throughout; weaker upward winds in TX. |
| Temperature PC1 | -2.45 | 0.65 | -3.76 | 4.3E-04 | Warmer temperatures throughout entire corridor |
| Temperature PC2 | 0.83 | 0.57 | 1.44 | 1.6E-01 | A stronger temperature gradient, cool to warm from MX to TX respectively. |
| Lag (29N to 30N) | 5.40 | 1.13 | 4.78 | 1.5E-05 |  |
| Lag (29N to 31N) | 8.09 | 2.33 | 3.47 | 1.0E-03 |  |
| Lag (29N to 32N) | 11.09 | 1.16 | 9.55 | 5.1E-13 |  |

**b) Parameter estimates selected for the "Combined" (push-pull-flow) top concept model (Table 1b) for summer arrival**
| <u>Top model parameters</u> | <u>Estimate</u> | <u>Std. Error</u> | <u>t-value</u> | <u>Pr(>\|t\|)</u> | <u>Parameter interpretation</u> |
| --- | --- | --- | --- | --- | --- |
| (Intercept) | 152.9 | 34.58 | 4.42 | 1.3E-04 |  |
| Lag (40N to 41N) | 6.0 | 1.85 | 3.26 | 2.9E-03 |  |
| Phase 2 slope model | -1.2 | 0.28 | -4.42 | 1.3E-04 | As predicted arrivals trend later, actual arrivals trend earlier relative to a 1:1 relationship |
| Temperature PC1 | -3.8 | 1.04 | -3.67 | 9.7E-04 | Warmer temperatures throughout entire corridor |
| Precipitation (Midwest) | -3.3 | 1.65 | -2.01 | 5.3E-02 |  |
| Greenup (TX) | 3.7 | 1.19 | 3.12 | 4.1E-03 |  |
| Winds PC2 | -2.3 | 1.25 | -1.84 | 7.7E-02 | TX winds blow more strongly westward and more weakly upward, but upward winds are stronger in the MW |
| Winds PC3 | 2.9 | 1.18 | 2.44 | 2.1E-02 | Upward winds are weaker, but eastward winds in the MW are stronger |
| Winds PC4 | -4.6 | 1.46 | -3.14 | 3.9E-03 | Northward and westward winds are stronger in TX, while northward and upward winds are weaker in the MW |
| Winds PC6 | 19.8 | 4.20 | 4.72 | 5.5E-05 | Northward winds are weaker throughout, while in TX, winds blow more strongly upward and westward. |

While typical arrival times have been known for at least 150 years^51^ (early to mid-March in spring and early to mid-May in summer), this study allowed us to narrow arrival window estimates from 11.4 to 6.6 days in spring and 16.6 to 9.4 days in summer (Table 1). These differences in estimated arrival dates can have substantial consequences for demographic research given the relatively short life spans of insects and the limited ability of neonates to withstand resource limitations^1,4,48,52^. Given this link between climate, migratory flow dynamics, and timing of milkweed emergence, we suggest that research on demographic consequences of limited milkweed during spring arrival should become a top priority.

An additional motivation for this study was to lay the groundwork for including migratory dynamics in projections of future monarch population trajectories, a factor that has not been included in research to date^53,54^. Our results show that the modeling framework and results we present are well-suited for this purpose. First, only environmental data are needed for model implementation, and this is necessary for its use in future projections based on climate. Further, the fit of both the spring and summer models provide for their use. This is true both of the correlative components fit by the GLM and the mechanistic model. In fact, this study allowed us to specifically evaluate the accuracy of our mechanistic model for FOY development time and then flight to the Midwest. Specifically, if the underlying mechanistic models were capturing actual development and flight rates, then we should see predicted FOY emergence times that occur, on average 17 days (our estimated flight time) before arrival and this is evident in Figure 3. This was also shown in the parameterization of the null summer arrival model, which should have a 1:1 slope between expected and observed average arrival dates. We assumed this 1:1 relationship with additional error being accounted for by the PPF variables, but we also included a slope modification parameter if there was bias in the estimates (see methods). Although the slope modification was a significant predictor in our final model (Table 2b), the slope was modified only slightly from 1 to 0.92. This means that both our estimates of development and flight times correctly reflect those occurring in the field, as suggested in Figure 3. The congruence between our null arrival model and observed average flight times provides evidence that the mechanistic modeling approach and parameters we used (T_max_, T_min_, k, and flight time) were both well supported. The use of hybrid correlational and mechanistic models for ecological projections has been suggested as an optimal approach because it can leverage the strengths of each^55^.

**Figure 3.**
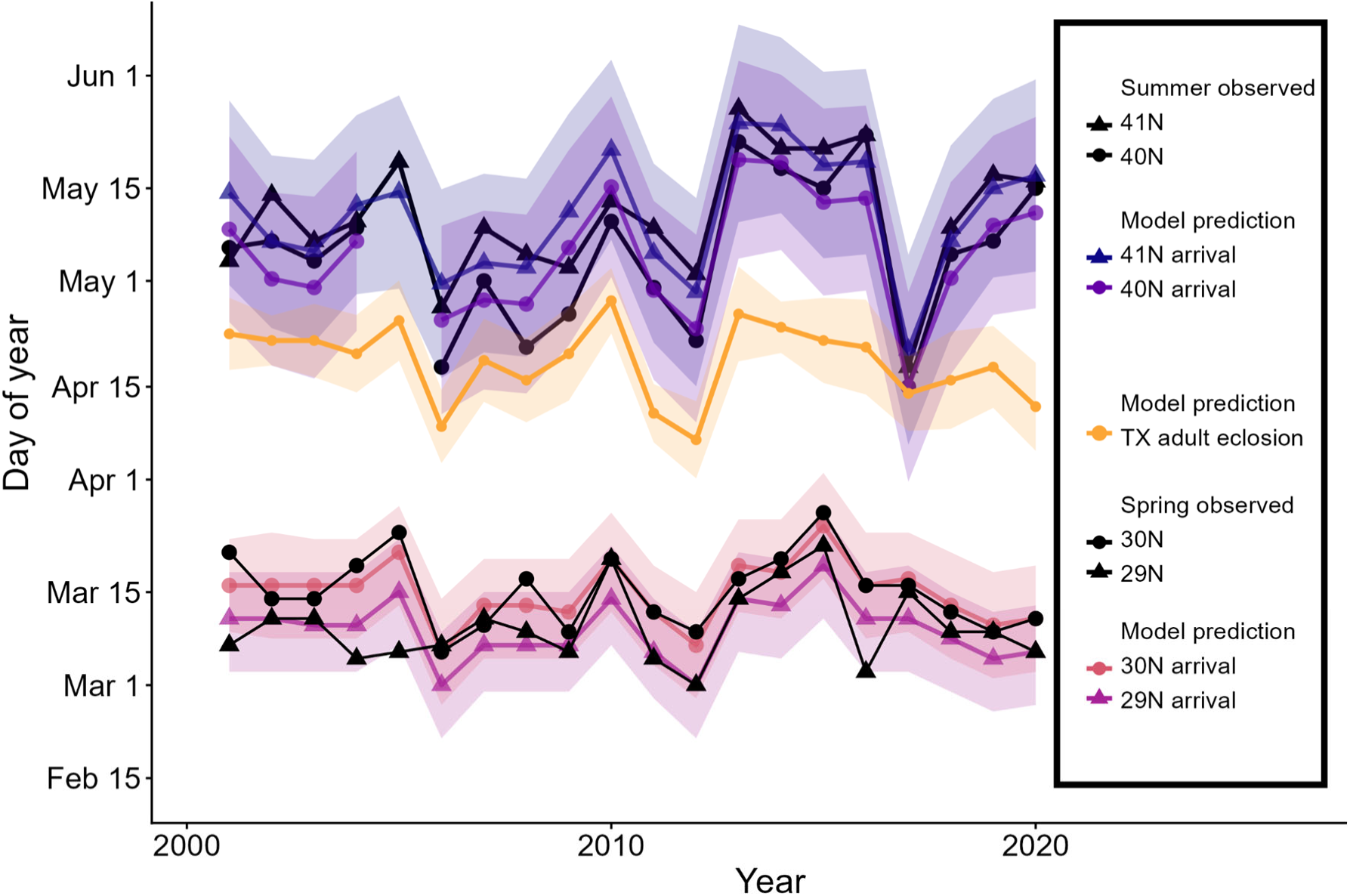
Model estimates (colored lines and bands) and observed timing (black lines) for monarch arrival timing based on top models (Table 2). Model predictions and confidence intervals are shown for spring arrival in 2 latitudinal bands in Texas (pinks) and earliest summer arrival in the US Midwest (blues). Arrival estimates from Journey North data are shown in black. For each arrival region, circles and diamonds distinguish separate latitudinal bands.

Climate patterns, and thus projections, for both spring and summer migration emerge from global circulation models (GCMs) that are substantially governed by the Hadley and Ferrel cells^56^. Therefore, projected covariates for the key weather conditions impacting monarch arrival timing at breeding grounds (temperature, rainfall, and wind) will be rooted in the strength and positioning of these cells. For the eastern migratory population of monarchs, Hadley dynamics appear to be the most important and so consideration of how this system may change in the future could be central to ecological projections of monarch populations. There is evidence that the northern edge of the Hadley cell has been shifting northward by about 0.05 degrees latitude per decade during spring, although there has been no change in intensity^56^. How these, or other changes in Hadley dynamics would translate into changes in spring arrival timing and location could be projected using our PPF model, especially if coupled with ecological projections of milkweed phenology and potential range-shifts. If a concomitant shift north in the spring range of monarchs were projected, this could be tested with JN data, although it may require a longer time-series than is currently available. Regardless, including migratory arrival timing could improve future-scenario projections of monarch dynamics, which already suggest declines will accelerate as emissions increase^53^, even without considering any potential mismatch as a component in the model.

Our results suggest potential demographic consequences of phenological mismatch in spring, but it was outside the scope of this study to determine if this dynamic is yet detectable on the ground. Current data density limitations meant our analysis was not sufficiently fine-grained to document how well monarchs may be able to locate the most abundant resources within their spring range, and thus ameliorate the potential demographic consequences of spring phenological mismatch. Fortunately, JN data (and those from similar platforms like iNaturalist.org^57^, and survey programs such as the Integrated Monarch Monitoring Program (IMMP)^58^) accumulate faster each year. Finally, new radio tags that can closely track detailed movement data across an individual’s migration has the substantial ability to transform how we study these phenomenon^59^ Combined with more sophisticated statistical models that can handle the sparse data associated with the monarch spring breeding season^40^ our ability to research large-scale, fine-grained demographic dynamics increases each year. These analyses could determine when (or if) lagging spatiotemporal distribution of milkweed resources impact FOY survivorship. We therefore suggest there is substantial potential to better grapple not only with how the eastern monarch may fare under a changing environment, but also the efficacy of the expanding milkweed-centered conservation efforts that are currently being implemented in hopes of reversing their long-term decline^60^.

## Acknowledgments

We are grateful to Elizabeth Howard for founding the Journey North program in 1994, and to Karen Oberhauser and the Monarch Joint Venture for their continued support curating the Journey North platform and providing open data access (available via http://journeynorth.org/monarch/). This work would not be possible without Journey North and its thousands of volunteers who have contributed observations on milkweed and monarchs over the 20 year study period. This paper was improved by the editing skills of Kate Epstein.

## Authors contributions

Conceptualization: NN

Methodology: NN, EAL

Investigation: NN, EAL, LR, RPG

Visualization: NN, EAL

Funding acquisition: NN, LR

Project administration: LR

Supervision: NN, LR

Writing – original draft: NN, LR

Writing – review & editing: NN, EAL, LR, RPG

## Data and materials availability

Code and processed data are available on github at (https://github.com/RiesLabGU/Neupane_Larsen_PushPullFlow_Monarch). Analyses used existing packages and programs, with minimal novel code.

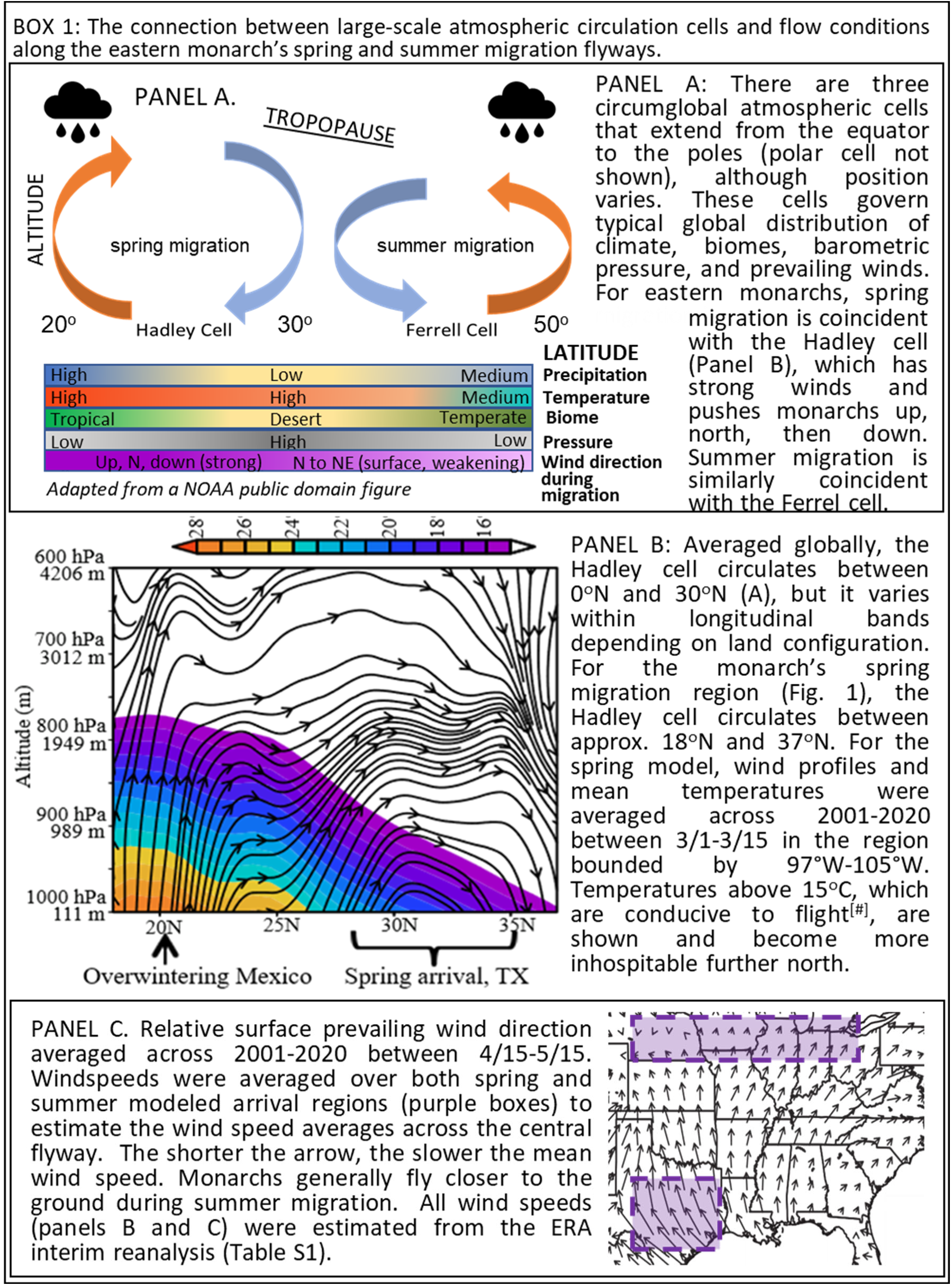

## Methods

### Monarch and milkweed phenology data

Phenology metrics were estimated from presence-only occurrence data for monarch adults and host plants that were obtained from Journey North (JN), a web-based community science portal focused on migratory species. One of the main benefits of using JN data for migratory research is that volunteers can specify if observations (eggs, adults and/or milkweeds for monarchs) are their “first sightings” of the season^1,2^. Thus, even though JN data are classified as “presence only” (a data type that usually can assume nothing about a reporter’s motivation for the data collected^3^), these data are particularly useful for calculating phenology metrics ^4^. We set 2001 as our start year, because it’s the first year of greenup data available (see below) and coincides with sufficient participation in JN to estimate phenology metrics (the program launched in 1996).

One of our main concerns with JN first sightings data was the potential for early monarch reports based on aberrant migratory behavior to bias our arrival estimates. In spring, our main concern was the small proportion of adults that overwinter in TX, mostly below 30°N^5^ and so we included spring butterfly observations only for days 50-150 (February 19-May 30/31) in each year. In summer, our main concern was individuals from Mexico overshooting the typical southern spring breed range and flying straight to the Midwest (when a lack of milkweed and cold temperatures make recruitment impossible). Thus, we restricted data extraction for summer arrival timing to days 50-150 (February 19-May 30/31) in each year. The spatial and temporal boundaries for JN downloads are illustrated in Figure 1 and detailed in Table S1.

To illustrate early arrival phenological patterns for monarchs and milkweed, we used JN data to plot the first 10 adult sightings and milkweed emergence from the spring and summer arrival zones, and these are shown in Fig. 2a . We refer to this set of sightings as our “first of the first” data set, because they are drawn from the subset of the JN data where volunteers are instructed to only report their first sighting from each season and then we include only the first 10 of these ”first sightings” for phenological estimates in each spatiotemporal unit. These “first of the first” sighting days were compiled into 1 degree latitudinal bands from 24-48°N (pooled across a longitudinal band from 106°W to 80°W longitude). This compilation region contains the majority of the central portion of their breeding range^6^. These sightings are overlaid on top of mean estimates of surface greenness using NDVI (the Normalized Difference Vegetation Index^7^) averaged across years from 2001-2020. We then used the same JN “first of the first” data to estimate mean timing of the progression of the northern moving front for both monarch arrival and milkweed emergence (Fig. 2b).

For the yearly metrics used in our predictive models, we calculated the arrival day of year (DOY) at the 15th percent quantile of JN sightings data using the quantileSE() function in the R broman package. JN sightings data were extracted from the same spatial boundaries used for environmental covariate data, detailed below. Monarch arrival estimates were calculated for one-degree latitudinal bands within each arrival region using JN monarch “first sightings” records within a season DOY window (DOYs 50-150 for spring arrival in Texas; DOYs 60-180 for summer arrival in the midwest). Because milkweed sightings are far scarcer than monarchs in JN, annual milkweed emergence metrics were calculated for the entire spring arrival region, rather than by latitudinal band, using all JN sightings of milkweed. Our use of the 15th quartile allows us to focus on the base of the upward portion of the arrival curve and thus disregard the earliest arrivals represented in the leading tail.

### Environmental covariate predictors

We used environmental covariates describing temperature, precipitation, wind, and greenness. To model our focal phases of the migratory cycle (spring and summer), we defined four regions for which we extracted covariate data. We extracted data from four regions, illustrated in Fig. 1.

1. The overwinter grounds in Mexico (MX), whose boundaries are well-known and located almost entirely within the boundaries or buffer zone of the Monarch Butterfly Biosphere Reserve^8^.
2. A focal zone within the spring arrival grounds in Texas (TX) (Fig. 1). This specific zone was chosen because it is in eastern TX, a primary arrival and spring breeding zone^9^, and it includes the dense observer populations in the cities of Dallas-Fort Worth, Houston, San Antonio and Austin. Thus, it represents the highest density of observations in eastern Texas.
3. The flight corridor between MX and TX. This zone was defined as a rectangle that begins just north of the mountain tops where overwintering occurs and extends to the southern portion of the US where they enter TX, but before they reach our spring breeding zone (Fig. 1).
4. An entry-point of the primary summer breeding grounds in the Midwest represented by a 2 degree latitudinal band^6^.

Note that we did not extract covariate data separately from the flight corridor between the TX and MW breeding grounds. This was because determining the best boundaries to characterize conditions in this more expansive migratory flight zone was not obvious as it was for spring. Further, preliminary explorations showed that conditions across the region were highly correlated regardless of different choices for regional zone boundaries. This is due to the fact that this zone is located within a single governing atmospheric cell (the Ferrel, see Box 1, Panel A) and that, unlike the spring migratory zone, the topographical profile between TX and the MW is relatively flat.

All weather variables were obtained from the North American Regional Reanalysis (NARR)^10^ and included temperature (T), wind (W) and precipitation (P). Wind variables were extracted along the two horizontal axes, North-South (N-S) and East-West (E-W), and the vertical Up-Down (U-D) axis. The combination of the three direction and speed metrics integrate to indicate the actual wind direction and speed. We show these averaged over 2001-2020 for the vertical and N-S horizontal axes (Box 1, Panel B, within the Hadley zone only) and also for the N-S and E-W horizontal axes across the monarch’s spring and summer breeding ranges (Box 1, Panel C). Temperature profiles illustrated along vertical clines in the Hadley zone (Box 1, Panel B). Two surface greenness metrics were used as a proxy that could represent cues for migrating monarchs of the densest availability of host plant or nectar resources. The first metric was mean surface greenness based on NDVI and the second was the date of midgreenup, using the Moderate Resolution Imaging Spectroradiometer (MODIS) Land Cover Dynamics Product C6 MCD12Q2^11^. This product provides data starting in 2001, which is one reason we set this year as the first for modeling. All candidate variables used, including spatial and temporal scope and grain, as well as data sources are detailed in Table S1.

The specific temporal and spatial scope for calculations of covariates for the spring and summer arrival models were partitioned as best as possible into departure (push), flight corridor (flow), and arrival (pull) zones. Since weather variables (and associated patterns such as plant growth) may be highly correlated due to synoptic climate dynamics^12^, we examined all predictor variables for collinearity both within and across zones. We found weak correlations with precipitation and greenness indices and so we were able to parse those into distinct push and/or pull variables in our candidate models (see VIF scores in Table 1). On the other hand, wind and temperature metrics were highly correlated and so we transformed these separately into principal components and the estimated loadings applied for each migratory phase are detailed in Table S2. Wind principle components (W.PCs) were considered to be “flow” variables only. For temperature, we used the regional temperature variables only for “push” or “pull” models. For models that would include temperature in both departure and arrival regions, we included temperature PC (T.PC) metrics

Table S2 details the T.PC and W.PC factor loadings calculated from the PCA analyses. We calculated temperature in the departure and arrival zones only and temperatures in the flyways were interpreted as a gradation between zones. T.PCs accounted for, respectively, 54 and 46% in spring variability and 83 and 17% of summer variability. Spring wind data included five variables: wind speeds in two directions at departure and arrival grounds (U-D and N-S, E-W winds showed very little variability in this zone, so were not included) and N-S winds only in the flight corridor. The resulting 5 PCs accounted for 57, 27, 8, 5, and 2% of variability. Summer wind data included six variables: wind speeds in all three directions (U-D, N-S and E-W) in departure and arrival zones only. The resulting 6 PCs accounted for 43, 25, 16, 10, 4, and 1% of variability.

### Push-pull-flow (PPF) arrival models

The PPF models for both spring arrival and summer arrival were estimated using linear models in the same 2-step approach. The first step was to develop a set of candidate models using different combinations of pull, push, and/or flow variables. We developed five candidate models: 1) pull only, 2) push only, 3) pull-push, 4) flow only, 5) all-combined (push, pull, and flow). We also developed a null model that included the number of JN observations as an offset variable and a distance lag. The distance lag was needed because each zone was separated into 1 degree latitudinal bands and arrival times are later for each successive band. In spring, the size of our focal zone spanned four bands that allowed us to use data from the four biggest cities (and so increased our sample size within the smallest area possible). This zone spanned 29°-33°N, so we included three lags: between the first (29-30°) and second (30-31°N) bands, the first and third bands (31-32°N) and the first and fourth bands (32-33°N). In summer, there are two bands (from 40°-42°N) so we included one lag between the first and second band. Each of the six candidate models (5 PPFs and 1 null) started as a full additive linear model with main effects only. We implemented a backwards stepwise procedure using AIC to remove candidate variables that had insufficient explanatory power. In the second step, we choose the best model from the six candidate models, also using AIC. The top model for spring is shown in Table 1a and the parameters estimates (with our interpretations of PC parameters) are shown in Table 2a. The arrival timing estimates from the spring model are then used in the first phase of our multi-level model for summer.

### Multi-level model for summer arrival

Although we used the same AIC model selection approach for both spring and summer models, the summer arrival model is more complex because it occurs in three distinct phases. The first phase is the date of spring arrival in Texas, and we use model-predicted (not estimates from sightings) dates from our spring arrival model (see above). Thus, our modeled estimate of spring arrival timing represents a sub-model for the second phase of the summer model. This phase estimates development time from egg, through larval and pupal stages culminating in emergence of FOY adults. We used a mechanistic model based solely on temperature and milkweed availability (detailed below). These phase-2 estimates were then used as a sub-model in the third phase, which used a fixed average flight time to the Midwest and then used the same AIC model selection approach as spring to model variability in observed summer arrival. Error is propagated across all three phases of the model. The restriction of using environmental variables only for phase 1 and 2 of the summer model means that it can be parameterized solely with environmental variables. This allows the model to be implemented in novel situations without on-the-ground monarch observations.

Our phase 2 mechanistic model of FOY emergence date included oviposition and development. Monarch oviposition requires both monarch arrival and milkweed availability, thus the predicted oviposition date was estimated as one day after predicted monarch arrival or, if monarchs’ estimated arrival occurred before milkweed estimated emergence (which happened in in three years in the southernmost latitudinal band), we instead used the estimated DOY of milkweed availability. Starting with the distribution of estimated oviposition dates, we modeled the development time from egg to adult using an established growing degree day (GDD) approach^13^. GDD models use daily temperature values to estimate how much development can occur each day based on mean temperature. Degrees are accumulated using minimum and maximum thresholds (T_min_ and T_max_) that are ideally determined in laboratory studies^13^. The date of developmental milestones (such as adult emergence) is then estimated by determining the total number of degree days needed to reach the specified milestone (k), a parameter that is also ideally measured in the lab^13^. Each day, the difference between the number of degrees between T_min_ and each day’s mean temperature (or T_max_, whichever is lowest) is calculated and degree days are accumulated until k is reached^13^. The GDD parameters we used in our model, 11.5C, 33C, and 325C for T_min_, T_max_, and k respectively, were reported (with errors) from laboratory experiments^14^.

We extracted daily maximum and minimum temperatures from the spring temperature data (Table S1) averaged across each latitudinal band within the region (29°N-33°N; 99°W-94°W) and calculated daily growing degree day values using a single-sine approximation. We produced a DOY estimate of adult emergence for monarch eggs laid in the spring region on the estimated arrival date from the phase 1 model plus estimated development time from phase 2. We calculated confidence intervals for adult emergence as a DOY range, running the deterministic GDD development model for a distribution of egg-laying DOYs and GDD requirements. For each year and latitudinal band, we simulated a distribution of adult emergence dates, beginning with 1000 DOYs from the distribution of predicted oviposition (egg-laying) dates. After excluding those before the estimated DOY of host plant availability, we sampled 1000 possible values of k (GDD requirement for development) from the distribution of GDD requirements in Zalucki^14^. For each predicted egg-laying date and each GDD requirement value, we calculated adult emergence DOY predictions by accumulating GDD until the given value of k was reached. The mean and standard deviation of the resulting emergence DOYs were used to describe the predicted adult emergence date for the generational development model. It is interesting to note that the generative FOY phase actually results in a small decrease in propagated error (from 3.89 to 3.02 days), because because on average, temperatures are increasing across the potential development period; thus, eggs laid earlier develop in colder temperatures (so develop relatively slowly), while development times are sped up when eggs are laid later in season and experience warmer temperatures. This range of dates then begins the clock for flight to the arrival zone in the Midwest (phase 3). In all years, the earliest adult emergence dates were predicted in the southernmost latitudinal band of the spring region.

Our final model component examined variation in flight times between the spring and summer regions relative to a null estimate in an average year. To set this null flight time, we used the estimated departure date from the southernmost latitudinal band (29°N-30°N) of the spring region to calculate expected average flight time to the summer region. Our estimated average flight time was 17 ± 1.4 days to travel from 29°N to 40°N (∼1223km), based on 71.6 km/day (range 66.6-78.1) km/day^15^. Flight speeds for fall have also been documented (7.5 +/-5km/hr^16^ and 7.27km/hr^17^). We assumed a 1:1 relationship between summer mean estimated and arrival dates for our model, but also a slope modification parameter to account for any bias. The top model for summer is shown in Table 1b and the parameters estimates (with our interpretations of PC parameters) for that model are shown in Table 2b.

